# Multimodal DTI-TMS assessment of the motor system in patients with chronic ischemic stroke

**DOI:** 10.1101/2019.12.24.886754

**Authors:** M. Nazarova, S. Kulikova, M. Piradov, A. Limonova, L. Dobrynina, R. Konovalov, P. Novikov, B. Sehm, A. Villringer, V. Nikulin

**Author notes:** Corresponding author: M. Nazarova, Centre for Cognition and Decision making, Institute for Cognitive Neuroscience, HSE University, Russia, Federal Center for Cerebrovascular Pathology and Stroke of the Ministry of Healthcare of the Russian Federation, Moscow, Russia. Address: Krivokolenny sidewalk 3, 101000, Moscow, Russia.

## Abstract

**Background and Purpose:** Despite the continuing efforts in multimodal assessment of the motor system after stroke, conclusive findings on the complementarity of functional and structural metrics of the corticospinal tract (CST) integrity and the role of the contralesional hemisphere are still missing. The aim of this work was to find the best combination of the motor system parameters, allowing classification of patients into three predefined groups of upper limb motor recovery.

**Methods:** 35 chronic ischemic stroke patients (47 [26–66] y.o., 29 [6–58] months post-stroke) with only supratentorial lesion and unilateral upper extremity weakness were enrolled. Patients were divided into three groups depending on the upper limb motor recovery. Non-parametric statistical tests and regression analysis were used to investigate the relationships among structural and functional motor system parameters, probed by diffusion tensor imaging (DTI) and transcranial magnetic stimulation (TMS). In addition, stratification rules were tested, using a decision tree classifier to identify parameters explaining motor recovery.

**Results:** Fractional anisotropy (FA) ratio in the internal capsule (IC) and absence/presence of motor evoked potentials (MEPs), were equally discriminative of the worst motor outcome group (96% accuracy). MEP presence diverged for two investigated hand muscles. Concurrently, for the three recovery groups’ classification, the best parameter combination was: IC FA ratio and Fréchet distance between the contralesional and ipsilesional CST FA profiles (91% accuracy). No other metrics had any additional value for patients’ classification.

**Conclusions:** This study demonstrates that IC FA ratio and MEPs absence are equally important markers for poor recovery. Importantly, we found that MEPs should be controlled in more than one hand muscle. Finally, we show that better separation between different motor recovery groups may be achieved when considering the whole CST FA profile.

## Introduction

Individual prognosis, with regard to upper limb motor function, largely depends on the integrity of the corticospinal tract (CST).^1^ There are two principally different approaches for CST integrity assessment: 1. structural - using T1, or diffusion-weighted MRI^1–3^ and 2. functional - using single pulse transcranial magnetic stimulation (TMS) based on probing motor evoked potentials (MEPs) from the affected hand.^4,5^ However, it is still unclear how much these approaches are interchangeable or complementary in different stroke periods.^4,6^ Despite being one of the most commonly used CST integrity parameters^3^, the method of detecting MEP presence is still not a unified one.^4,5, 7–11^ It is still not clear, whether it is necessary to study more than one muscle and which hand muscles are the most informative.^12–15^ Another possible important selection biomarker, for rehabilitation intervention choice, is the state of the contralesional motor areas and the excitability balance between the hemispheres.^16,17^ It is becoming clear that popular inhibitory stimulation of the contralesional primary motor cortex, is not effective when applied in a “one-size-fits-all” way.^17–19^

In this study, we combined MRI- and TMS-based measures of the motor system in a cohort of chronic supratentorial ischemic stroke patients. Specifically, we aimed at classifying patients into three predefined groups of the upper limb motor outcome. We investigated: (1) whether the use of two, and not one, hand muscles has an added value for MEPs detection; (2) whether structural and functional CST metrics are complementary to each other; and (3) whether additional metrics, such as MEPs parameters, evoked by single and paired pulse TMS (ppTMS) of any of the hemispheres, and corpus callosum (CC) structural integrity, can improve the classification, based solely on the CST.

## Materials and Methods

The data that support the findings of this study are available from the corresponding author on reasonable request.

### Population

Thirty-five patients (13 women) aged from 26 to 66 y.o. (mean age 46.6 ± 10.1 y.o.) were enrolled, 28 of them underwent both DTI and TMS investigations (Table 1/Table I in the online-only Data Supplement). Inclusion criteria were: chronic supratentorial ischemic stroke, leading to the upper limb paresis in the acute period of stroke. Time after stroke was 6 months and more (29.1±21.8 months). MRI confirmed a diagnosis of only ischemic lesion. Exclusion criteria included: neurological conditions, e.g. signs of the small vessel disease rated Fazekas score>1, any standard contraindications for MRI and/or TMS, pregnancy, any serious somatic issues, Montreal Cognitive Assessment score < 25. Strict exclusion criteria, regarding white matter lesions, led to mostly young and middle age of the investigated patients. This study was approved by the IRB of the Research Center of Neurology N312, a written informed consent was obtained from all patients.

**Table 1.** Distribution of the patients in three groups of motor outcome

|  | DTI | DTI+TMS | TMS |
| --- | --- | --- | --- |
| Total number | 32 | 28 | 31 |
| Mean age ± sd | 45.9 ± 9.7 | 46.6 ± 9.96 | 47.3 ± 10.3 |
| men : women | 19 : 13 | 16 : 11 | 18 : 13 |
| Group 1 - *good* | 8 | 7 | 9 |
| Group 2 - *moderate* | 6 | 6 | 7 |
| Group 3 - *bad* | 18 | 14 | 15 |

### Behavioral assessment

Hand function was assessed using the Fugl-Meyer Assessment scale for upper limb (FM-UE) and a scale that is commonly used in Russia - spastic paresis scale of Institute of Neurology (SPS_IN*, 1982,\*\**)^20^ (online-only Data Supplement). Based on these scales, patients were divided into three outcome groups: 1 - good (1, 2 points in SPS_IN, active use of the hand in everyday life with different levels of dexterity, FM-UE >55 scores); 2 - moderate (3 points SPS_IN, limited use of the hand in everyday life, FM-UE 30-55); 3 - bad hand motor recovery (4, 5 points SPS_IN, no use of the hand in everyday life, FM-UE<30).

### MRI investigation

MRI data was acquired using a 1.5T Magnetom Avanto scanner (Siemens, Germany). Structural T1-weighted images (MPRAGE, 1 mm isotropic voxel; acquisition matrix 256×256) were used for MRI-guided TMS. Diffusion-weighted data were acquired using 20 gradient directions, b-value=1000 s/ mm^2^ and voxel size 1.8x 1.8x 5.0 mm, following the routine clinical protocol.

### TMS investigation

MRI-navigated TMS was performed using a Nexstim eXimia stimulator with TMS-compatible eXimia electromyography (EMG). MEPs were recorded by placing 0.6-cm^2^ surface EMG electrodes over two hand muscles - abductor pollicis brevis (APB), and extensor digitorum communis (EDC).^21^

Single pulse TMS of both hemispheres was performed using figure-of-eight biphasic coil (version 3.2.2, Focal Bipulse, outer winding diameter 70 mm). First, “rough mapping”^22^ starting from the “motor knob” region^23^ in the primary motor cortex and proceeding with somatosensory and premotor areas was performed to find optimal “hotspots” for both investigated muscles (similar to Krieg et al.^22^). Once both hotspots were found, the resting motor threshold (RMT) was determined for each muscle, defined as the lowest intensity sufficient to evoke MEP with the amplitude of 50 µV in a resting muscle in about 5 out of 10 stimuli.^24^ RMT was estimated as a percentage of the maximum intensity of the stimulator. In patients whose RMT for an investigated muscle (separately for APB and EDC) was higher than 99% of the maximal stimulator output, the criterion was changed to eliciting MEP with 100% intensity with preactivation. MEP was considered as present (MEP+) when it was possible to obtain MEP higher than 50 µV in any of the tested muscle with a consistent latency in 4 out of 8 consecutive trials during rest or attempted/imagined contraction of the affected and/or contralesional hand, which is similar to the approach described in the PREP2 algorithm ^7^.

The ppTMS protocol is described in detail in the online-only Data Supplement.

## Data analysis

### DTI data

DTI data were corrected for artifacts as described in.^25^ FA maps were calculated using BrainVISA^26,27^ and normalized in SPM8 (Welcome Trust Centre of Neuroimaging, UK). CС and CST masks were manually delineated on averaged normalized FA maps using BrainVISA^26^ from an additional group of 30 healthy subjects from a separate study (18 women, mean age 46.9 ± 16.5 y.o.) with the same acquisition parameters.^28^ Images of the right-hemispheric stroke patients were flipped for further comparison in SPM8, using MRIcron (http://www.nitrc.org/projects/mricron). In the CST, we manually defined regions of interest (ROIs) in the posterior limb of the IC and in the cerebral peduncle (CP). FA asymmetry was calculated as the ratio between ipsilesional and contralesional FA. Centroids for the CST from Connectomist^27^ were projected to the subjects’ data to measure FA values at 30 equidistant points. To assess the similarity between ipsi- and contralesional FA CST profiles, we used Fréchet distance, that takes into account the location and ordering of the points along the curves.^29^ Formally, the Fréchet distance *d_f_*(*A, B*) between curves A and B is defined as follows:

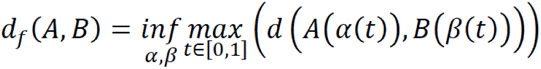 where *α*(*t*), *β*(*t*) are any continuous non-decreasing parameterizations of the curves A and B.^30^ To our knowledge, it is the first time Fréchet distance is applied for analysis of ipsi- and contralesional FA CST profiles.

There is currently no consensus which CC areas are the most discriminative for stroke patients with motor deficit.^31–33^ Thus, we decided to perform voxel-based comparison of FA maps between the best (Group1) and the worst recovered patients (Group3). Normalized FA maps of Group3 and Group1 were compared using two-sample t-test with the CC masks using SPM8. Voxels were identified as significant if p_uncor ≤ 0.01 on the cluster level, family-wise error corrected. For data visualization, we utilized xjView.814 (https://www.alivelearn.net/xjview). FA in this CC area was used for further analysis.

### TMS data

The MEPs peak-to-peak amplitudes were calculated online using the eXimia software. During the initial preprocessing, EMG data were visually inspected for artifacts. Trials with the pre-stimulus noise larger than 50 µV were excluded.^22^ We also estimated RMT ratio: RMT ipsilesional /RMT contralesional hemisphere. In contrast to Kemlin et al.^34^, we did not assign 110% of RMT in cases, when MEPs were absent in the contralateral muscles (see the discussion).

### Patients’ classification regarding upper extremity motor outcome

Classification of patients was performed using WEKA 3.8.^35^ Among potential characteristic features, we considered all the investigated DTI and TMS metrics: FA asymmetry in the СST ROIs (FA IC, FA CP), FA asymmetry in 30 pairs of symmetric points along the CST FA profile, Fréchet distances between the CST FA profiles^29^, FA in the CC ROI, RMTs and MEPs presence for ABP and EDC, ppTMS metrics in APB in both hemispheres.

We performed feature selection, using ClassifierSubsetEval function in WEKA 3.8^35^, evaluating attribute subsets by measuring the information gain with respect to each of the three classes. This procedure retained the following features: FA IC, FA asymmetry at positions x10-x12 along the CST (approximately at the level of the delineated IC), and Fréchet distances between CST FA profiles.

For classification, we used decision tree J48 with 10-fold cross-validation. Decision trees are predictive models, allowing mapping observations of item characteristics (DTI and TMS metrics) to the target values (motor outcome). Here decisions have a tree-like structure (Figure I in the online-only Data Supplement) with nodes for testing features, and leaves, each labeled with a unique class name. Splits in J48 classifiers are based on individual attributes, so we additionally considered linear combinations of up to three attributes.

### Statistical analysis

Apart from the procedures described above, further statistical analysis was performed using GraphPad Prism. We used Spearman’s correlation between TMS and DTI metrics. Mann-Whitney U-Test was applied for comparing FA metrics, RMTs, and ppTMS phenomena among the recovery groups and for comparing DTI parameters between MEP+ and MEP- patients. False discovery rate (FDR) correction was performed to control for multiple comparisons in Matlab (Natick, USA). Confidence intervals for Mann-Whitney U-test were calculated as in Perme et al.^36^ For the Spearman rank test, confidence intervals were calculated as in Bonett et al.^37^

## Results

### Structural DTI assessment of CST and CC integrity

*Corticospinal tract:* FA ratio in IC and CP showed significant differences between Group3 and the two other groups (Figure 1A, B). While Fréchet distance between ipsi- (Figure 1C) and contralesional (Figure II in the online-only Data Supplement) CST FA profiles were significantly different among all the groups (Mann-Whitney U-test) (Figure 1D).

**Figure 1.**
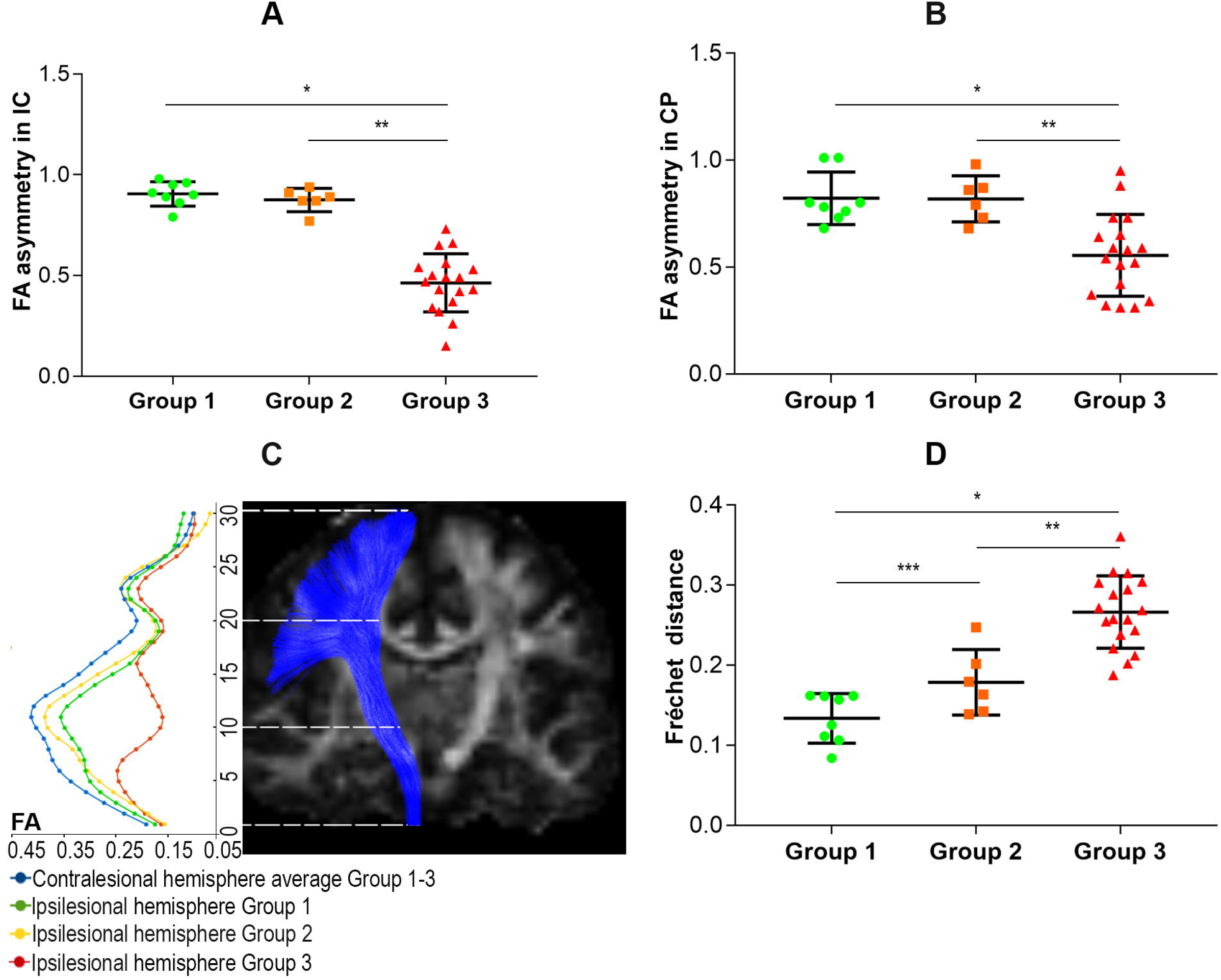
Structural assessment of CST in three groups of motor outcome. (A) FA ratio in the internal capsule (IC) (Mann-Whitney U-test: *θ = 1.0 [95% CI, 0.88 to 1.00], *p= 0.0002; **θ = 1.00 [95% CI, 0.89 to 1.00], **p= 0.0007) and (B) cerebral peduncle (CP) – significant difference between Group 3 and two other groups, Group 3 can be delineated from two other groups only using IC FA (Mann-Whitney U-test: *θ = 0.90 [95% CI, 0.62 to 0.98], *p= 0.005; **θ = 0.88 [95% CI, 0.61 to 0.97], **p= 0.0137); (C) FA profiles of the ipsilesional CST; (D) Fréchet distance between ipsi- and contralesional FA CST profiles - significant difference among all groups of motor recovery (Mann-Whitney U-test: *θ = 1.0 [95% CI, 0.90 to 1.0], *p= .0002; **θ = 0.94 [95% CI, 0.59 to 0.99], **p=0.0039; ***θ = 0.83 [95% CI, 0.46 to 0.97], ***p= 0.0426).

*Corpus callosum:* the most diverging CC area between well and poorly recovered patients overlapped by 76% with the CC hand motor area from the Brainnetome atlas (Figure 2A). FA values there were significantly different between Group1 and Group3 (Figure 2B) and correlated significantly with FA in the ipsilesional (Figure 2C), but not in the contralesional IC (Figure 2D).

**Figure 2.**
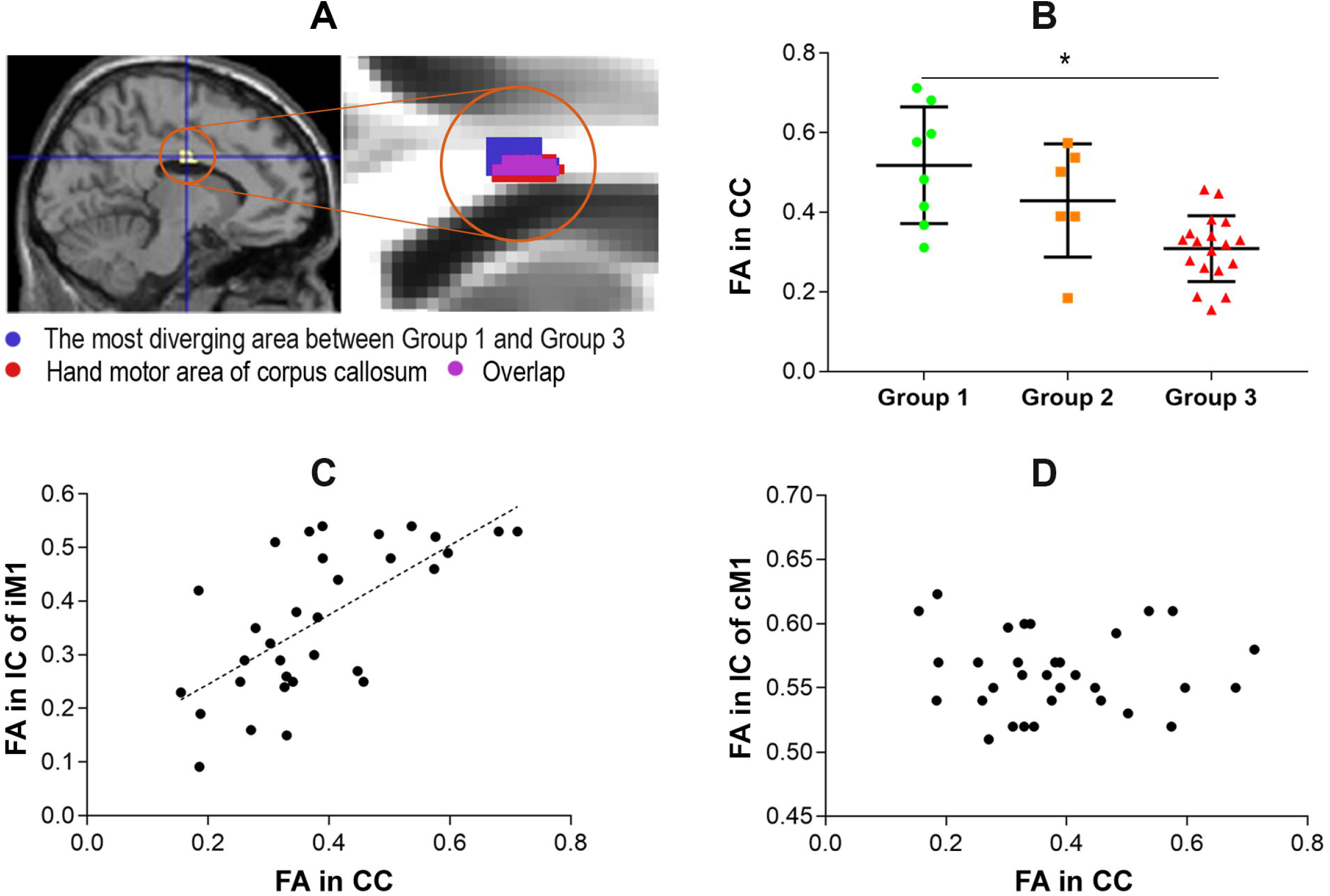
(A) Voxel-based comparison of the FA in the corpus callosum (CC) between groups 1 and 3 and zoomed-in version of this ROI (dark blue) and its overlap (magenta) with CC hand motor region from the Brainnetome atlas (red); (B) Comparison of FA values in the selected CC ROI among three groups of motor recovery – significant difference between groups 1 and 3 (Mann-Whitney U-test: *θ = 0.89 [95% CI, 0.55 to 0.98], *p=0.0061); (C) Correlation of FA in the CC ROI with FA in the internal capsule (IC) of the ipsilesional hemisphere (iM1) (Spearman correlation coefficient = 0.67, p<0.0001, [95% CI, 0.40 to 0.83]) and (D) no correlation of CC FA values with the IC FA values in the contralesional hemisphere (cM1).

### Functional TMS assessment of the ipsilesional hemisphere

#### Single pulse TMS

All Group1 patients were MEP+ for both muscles of the affected hand. All Group2 patients were also MEP+, however, in three patients, MEPs were obtained only from a single muscle, in two patients only from APB (RMTs were 98% and 64%, respectively); in one - from EDC (RMT was 46%). RMT had a tendency to be higher in Group2, but it did not differ significantly from that in Group1. In Group3, only one patient was MEP+. Importantly, his RMTs were lower than the average RMT for Group1. Interestingly, this patient was among the four patients with the shortest time after stroke (6 months). As a normalized version of the affected side RMT, we calculated RMT ratio between ipsilesional and contralesional RMT only for those muscle pairs with MEPs. No significant difference for either APB or EDC muscles was found after FDR correction; also, there is an apparent tendency towards higher values in Group2 (Table 2).

**Table 2.** Mean RMT, RMT ratio and MEP+/- for the ipsilesional hemisphere for three groups of motor outcome

|  | Group 1 | Group 2 | Group 3 |
| --- | --- | --- | --- |
| MEP+ | 9 | 7 (in 3 – in the only muscle) | 1 |
| MEP- | 0 | 0 | 14 |
| RMT APB,<br>mean ± sd | 46.56±16.43 | 65.33±22.51 | - |
| RMT ratio APB,<br>mean ± sd | 1.08±0.27 | 1.42±0.35 | - |
| RMT EDC,<br>mean ± sd | 47.44±16.33 | 63.80±24.45 | - |
| RMT ratio EDC,<br>mean ± sd | 1.09±0.32 | 1.51±0.20 | - |

#### SICI and ICF phenomena in stroke

No significant difference for SICI and ICF phenomena in the ipsilesional hemisphere (only in MEP+ patients) was found. An inversed SICI phenomenon in the ipsilesional primary motor cortex (SICI/SP>1) was found in three patients: in one Group1 patient and in two Group2 patients. No between-group difference was observed for the ICF.

### Functional TMS assessment of the contralesional hemisphere

#### Single pulse TMS

RMT from the contralesional primary motor cortex did not differ among the groups (Figure 3A). Notably, RMT of the contralesional and ipsilesional primary motor cortex were highly correlated in MEP+ patients (Figure 3B).

**Figure 3.**
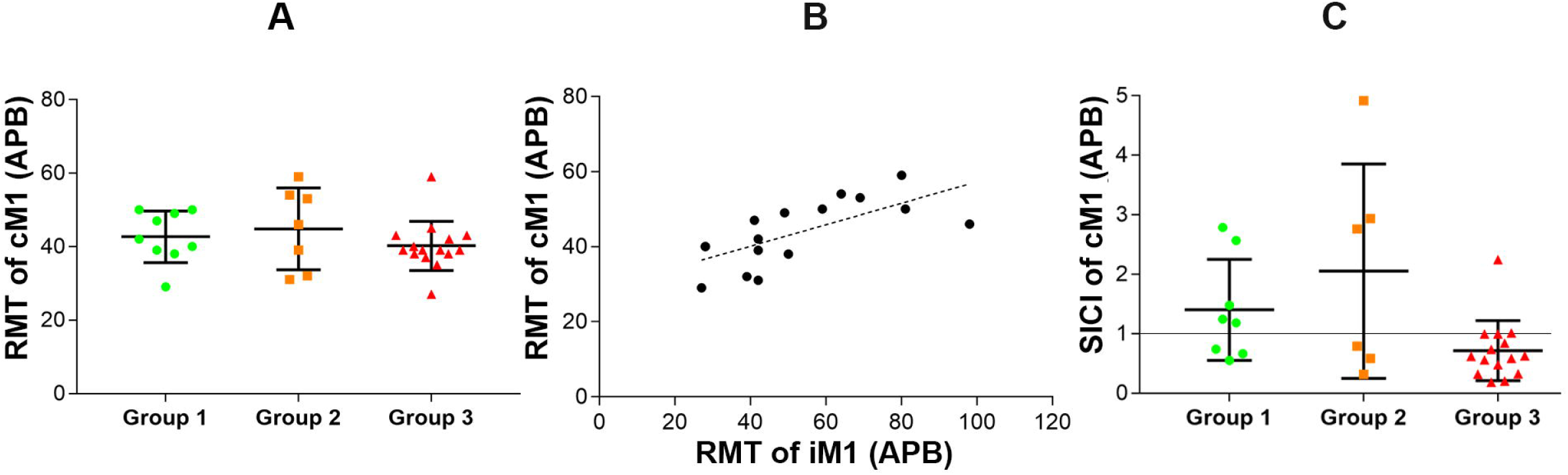
Contralesional hemisphere TMS assessment. (A) No difference among the recovery groups for RMT of the contralesional hemisphere for the APB; (B) Correlation between RMT of the contralesional hemisphere (cM1) and the ipsilesional hemisphere (iM1) in MEP+ patients (Spearman correlation coefficient = 0.76, p<0.001, [95% CI, 0.44 to 0.91]); (C) Short interval intracortical inhibition (SICI) in the contralesional hemisphere: lack of inhibition may be observed in some patients from good and moderate recovery groups, but not in poorly recovered patients, though the difference between Group 1 and 3 is not significant (Mann-Whitney U-test, p= 0.15).

#### ppTMS

Inhibition (SICI/SP<1) in the contralesional hemisphere was found in all Group3 patients. The SICI phenomenon was inverted (SICI/SP>1) in the majority of Group1 (5 out of 8) and Group2 patients (3 out of 6). Groups1 and 3 were significantly different. Disinhibition in the contralesional hemisphere was more characteristic for Group1 (Figure 3C). No significant difference of the ICF phenomena in the contralesional hemisphere was found.

### Association between structural and functional assessments of the CST integrity

Since MEP presence was the best functional discriminating factor, presumably reflecting the CST integrity, we used it to discriminate structural parameters. In all cases structural parameters for MEP+ and MEP- groups were significantly different: FA asymmetry in the IC and CP, Fréchet distance, and FA in the CC ROI. The best separation between MEP+ and MEP- groups was observed for IC FA ratio (Figure III A-D in the online-only Data Supplement).

ppTMS phenomena (SICI, ICF), probed in the contralesional primary motor cortex, did not correlate with the CC structural integrity.

### Classification of patients to motor outcome groups using a decision tree classifier

Both IC FA asymmetry and MEP presence were equally discriminative for Group3 versus the two other groups (96% accuracy). When other parameters such as CST, and the CC integrity, were added, their best predicting combination included two CST integrity parameters, FA IC asymmetry, and Fréchet distance (3 classes, 91% accuracy) (Figure 4, Figure IV in the online-only Data Supplement). No other metrics had any additional value for patients’ classification.

**Figure 4.**
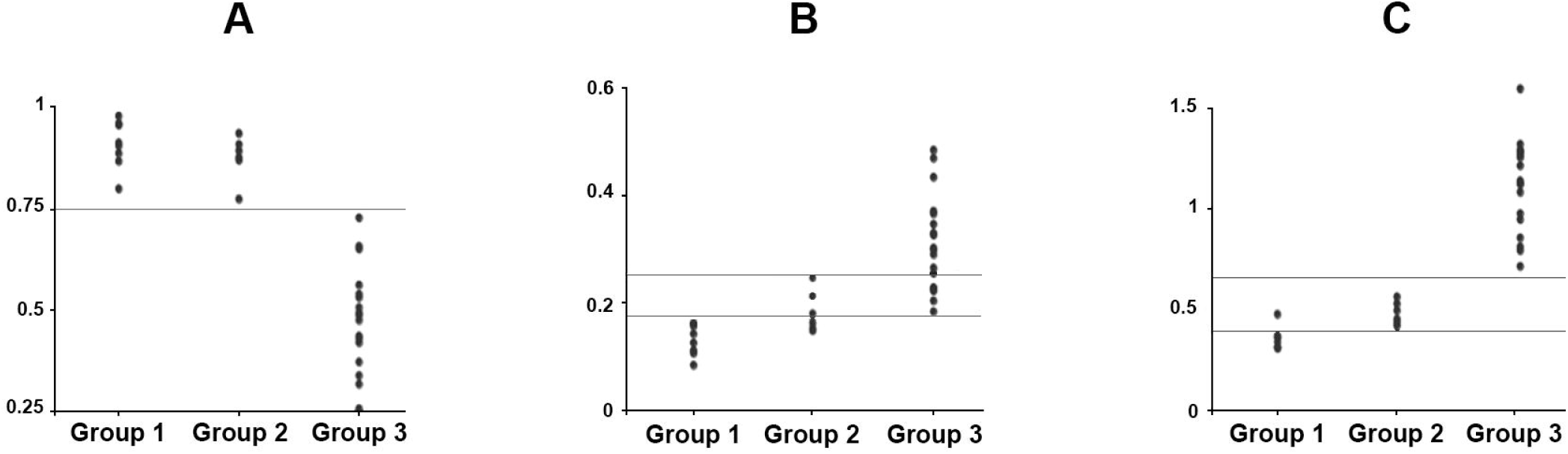
Classification of patients using CST integrity metrics. (A) FA asymmetry in the IC separates Group 3 from two other groups with 96% accuracy; (B) Fréchet distance between ipsi- and contralesional CST FA profiles differentiates better groups 1 and 2; (C) the combination of these two measures is able to separate all three groups from each other with 91% accuracy.

## Discussion

We probed the relationships between upper limb motor deficit in chronic supratentorial ischemic stroke patients, and several structural-functional parameters of the motor system. We aimed at investigating whether factors, apart from the CST integrity, would have an additional value for patients’ classification.

### Structural CST integrity for classification

Previous studies suggested that IC FA values provided an excellent separation of the group with bad motor recovery.^38,39^ However, no or little difference was found between motor recovery groups when comparing FA over the CST, as in the recent study by Kemlin and colleagues.^34^ Such a result might stem from the lack of ROIs specificity. We considered both ROI-analysis and FA profiles along the CST. Apart from replicating the finding regarding the IC FA^38,39^, we also showed that the combination of IC FA asymmetry and Fréchet distance is superior in classifying motor-outcome groups compared to CST ROI-based approaches.

### MEP presence for classification

MEPs presence had the same classification value as IC FA ratio. Importantly, in three out of seven Group2 patients, MEPs were present in a single muscle which differed among patients. It is worth noting that such “single MEP+ muscle” situation happened only in the Group2. Therefore, we assume that it may be informative to probe MEPs in several muscles.

### Resting motor threshold in the ipsilesional hemisphere

We suggest that such a multi-muscle approach might also be applicable for other TMS parameters, such as RMT. Indeed, we showed that RMT, in the ipsilesional hemisphere, may vary among different muscles. When performing between-group comparisons, no significant difference between Group1 and Group2 for RMTs or RMTs ratio was found. However, this might be because of a small number of MEP+ patients in Group2. Indeed, in a recent study^34^, the RMT ratio was the only metric correlating with the motor performance. The authors proposed to assign 110% RMT to the affected side for the most severely affected MEP- patients.^34^ However, we decided to avoid this. We suppose that RMT ratio does not reflect an interhemispheric balance, but may be rather considered as a normalized value of the ipsilesional RMT. As it was shown previously^12^, and in our study, contralesional RMT does not change after stroke and, thus, does not differ among patients with different recovery level. At the same time, when assigning 110% RMT to an MEP- hand, such ratios would reflect only contralesional RMT, which are likely to relate to its premorbid value. Thus, we suppose that RMT ratio should be considered only for MEP+ patients, i.e. patients with moderate-to-mild deficit.

### Resting motor threshold in the contralesional hemisphere

Contralesional RMTs were similar among the recovery groups and RMTs from both sides correlated strongly in MEP+ patients. For the contralesional CST, the correlation of RMT and FA was weak but significant, similar to the previous findings in healthy subjects.^15^ Interestingly, when investigating ipsilesional FA IC and contralesional RMTs, two segregated clusters appeared with a significant negative correlation between FA IC and RMT only in the sub-population of patients with moderate and good outcome (FA IC>0.4). A possible interpretation of these results is that RMT is associated, at least to some extent, with the CST integrity. However, such association did not seem to be more prominent in the ipsilesional compared to the contralesional side.

### Corpus callosum integrity

It has been suggested that CC integrity may serve as a biomarker in the chronic stroke.^31^ However, previous data on the most relevant CC regions, and on their contribution depending on paresis severity, are contradictory. Indeed, one study suggested that FA in the CC motor region, explains the motor deficit scores better than FA in the CST in a subgroup of patients with minor motor impairment.^33^ While in other studies, relationships between CC integrity and recovery were shown for the prefrontal CC region^31,40^, and in one of them^31^, such association existed only in a subgroup of the most severely affected patients. We defined the CC ROI based on voxel-based comparison of the “good” and “bad outcome” groups. Interestingly, the resulting ROI overlapped by 76% with the CC hand motor region from the Brainnetome atlas.^41^ However, FA in this region did not have additional value for classification. It is not surprising, considering that FA in this region is highly correlated with the ipsilesional IC FA. Conversely, no correlation with the contralesional CST was observed which is consistent with the previous findings in healthy volunteers.^42^

## Conclusions

We replicated the finding that IC FA asymmetry and MEPs absence can serve as markers for the poorest hand motor recovery with a similar predictive value. As a novel finding, we demonstrated an additional value of the difference between ipsi- and contralesional CST FA profiles. Furthermore, we showed for the first time that MEPs should be controlled in more than one hand muscle. MEPs in just one of the investigated muscles were observed only in patients with moderate recovery, highlighting the possibility of false negative MEPs feature in this group.

Other principal findings are related to the interhemispheric interactions: (1) the most prominent difference between well- and poorly-recovered patients in the CC, is located primarily in the CC hand motor region; (2) the state of the contralesional motor cortex is not associated with CC integrity.

## References

1. Rondina JM, Park CH, Ward NS. Brain regions important for recovery after severe post-stroke upper limb paresis. J Neurol Neurosurg Psychiatry. 2017;88:737–743.

2. Kim B, Fisher BE, Schweighofer N, Leahy RM, Haldar JP, Choi S, Kay DB, Gordon J, Winstein CJ. A Comparison of Seven Different DTI-derived Estimates of Corticospinal Tract Structural Characteristics in Chronic Stroke Survivors HHS Public Access. J Neurosci Methods. 2018;304:66–75.

3. Boyd LA, Hayward KS, Ward NS, Stinear CM, Rosso C, Fisher RJ, Carter AR, Leff AP, Copland DA, Carey LM, Cohen LG, Basso DM, Maguire JM, Cramer SC. Biomarkers of Stroke Recovery: Consensus-Based Core Recommendations from the Stroke Recovery and Rehabilitation Roundtable*. Neurorehabil Neural Repair. 2017;31:864–876.

4. Stinear CM, Barber PA, Coxon JP, Byblow WD, Fleming MK, Smale PR. Functional potential in chronic stroke patients depends on corticospinal tract integrity. Brain. 2006;130:170–180.

5. Stinear CM, Barber PA, Petoe M, Anwar S, Byblow WD. The PREP algorithm predicts potential for upper limb recovery after stroke. Brain. 2012;135:2527–2535.

6. Hayward KS, Lohse KR, Bernhardt J, Lang CE, Boyd LA. Characterising Arm Recovery in People with Severe Stroke (CARPSS): Protocol for a 12-month observational study of clinical, neuroimaging and neurophysiological biomarkers. BMJ Open. 2018;8:26435.

7. Stinear CM, Byblow WD, Ackerley SJ, Smith MC, Borges VM, Barber PA. PREP2: A biomarker-based algorithm for predicting upper limb function after stroke. Ann Clin Transl Neurol. 2017;4:811–820.

8. Lee JW, Kwon YM, Jang SH. Predictability of Motor Outcome According to the Time of Motor Evoked Potentials From the Onset of Stroke in Patients With Putaminal Hemorrhage. Ann Rehabil Med Orig Artic Ann Rehabil Med. 2015;39. doi:10.5535/arm.2015.39.4.553.

9. Yarossi M, Patel J, Qiu Q, Massood S, Fluet G, Merians A, Adamovich S, Tunik E. The Association Between Reorganization of Bilateral M1 Topography and Function in Response to Early Intensive Hand Focused Upper Limb Rehabilitation Following Stroke Is Dependent on Ipsilesional Corticospinal Tract Integrity. Front Neurol. 2019;10:258.

10. Hoonhorst MHJ, Nijland RHM, van den Berg PJS, Emmelot CH, Kollen BJ, Kwakkel G. Does Transcranial Magnetic Stimulation Have an Added Value to Clinical Assessment in Predicting Upper-Limb Function Very Early After Severe Stroke? Neurorehabil Neural Repair. 2018;32:682–690.

11. Schambra HM, Xu J, Branscheidt M, Lindquist M, Uddin J, Steiner L, Hertler B, Kim N, Berard J, Harran MD, Cortes JC, Kitago T, Luft A, Krakauer JW, Celnik PA. Differential Poststroke Motor Recovery in an Arm Versus Hand Muscle in the Absence of Motor Evoked Potentials. Neurorehabil Neural Repair. 2019. doi:10.1177/1545968319850138.

12. Rosso C, Lamy JC. Does resting motor threshold predict motor hand recovery after stroke? Front. Neurol.. 2018;9:1020.

13. Herbsman T, Forster L, Molnar C, Dougherty R, Christie D, Koola J, Ramsey D, Morgan PS, Bohning DE, George MS, Nahas Z. Motor threshold in transcranial magnetic stimulation: The impact of white matter fiber orientation and skull-to-cortex distance. Hum Brain Mapp. 2009;30:2044–2055.

14. Hübers A, Klein JC, Kang JS, Hilker R, Ziemann U. The relationship between TMS measures of functional properties and DTI measures of microstructure of the corticospinal tract. Brain Stimul. 2012;5:297–304.

15. Klöppel S, Bäumer T, Kroeger J, Koch MA, Büchel C, Münchau A, Siebner HR. The cortical motor threshold reflects microstructural properties of cerebral white matter. Neuroimage. 2008;40:1782–1791.

16. Pino G Di, Pellegrino G, Assenza G, Capone F, Ferreri F, Formica D, Ranieri F, Tombini M, Ziemann U, Rothwell JC. Modulation of brain plasticity in stroke : a novel model for neurorehabilitation. Nat Publ Gr. 2014;10:597–608.

17. Ovadia-Caro S, Khalil AA, Sehm B, Villringer A, Nikulin V V., Nazarova M. Predicting the response to non-invasive brain stimulation in stroke. Front Neurol. 2019;in press:302.

18. Harvey RL, Edwards D, Dunning K, Fregni F, Stein J, Laine J, Rogers LM, Vox F, Durand-Sanchez A, Bockbrader M, Goldstein LB, Francisco GE, Kinney CL, Liu CY, NICHE Trial Investigators *. Randomized Sham-Controlled Trial of Navigated Repetitive Transcranial Magnetic Stimulation for Motor Recovery in Stroke. Stroke. 2018;49:2138–2146.

19. Xu J, Branscheidt M, Schambra H, Steiner L, Widmer M, Diedrichsen J, Goldsmith J, Lindquist M, Kitago T, Luft AR, Krakauer JW, Celnik PA, Kim N, Harran MD, Hertler B, Cortes JC. Rethinking interhemispheric imbalance as a target for stroke neurorehabilitation. Ann Neurol. 2019;85:502–513.

20. Stoliarova LG, Kadykov AS TG. Scoring system for the status of motor functions in patients with post-stroke paralyses. Zh Nevropatol Psikhiatr Im S S Korsakova. 1982;82:15–8.

21. Selvanayagam VS, Riek S, Carroll TJ. A systematic method to quantify the presence of cross-talk in stimulus-evoked EMG responses: Implications for TMS studies. J Appl Physiol. 2012;112:259–265.

22. Krieg SM, Lioumis P, Mäkelä JP, Wilenius J, Karhu J, Hannula H, Savolainen P, Lucas CW, Seidel K, Laakso A, Islam M, Vaalto S, Lehtinen H, Vitikainen AM, Tarapore PE, Picht T. Protocol for motor and language mapping by navigated TMS in patients and healthy volunteers; workshop report. Acta Neurochir (Wien). 2017;159:1187–1195.

23. Yousry TA, Schmid UD, Alkadhi H, Schmidt D, Peraud A, Buettner A, Winkler P. Localization of the motor hand area to a knob on the precentral gyrus. A new landmark. Brain. 1997;120 (Pt 1:141–57.

24. Rossini PM, Barker AT, Berardelli A, Caramia MD, Caruso G, Cracco RQ, Dimitrijević MR, Hallett M, Katayama Y, Lücking CH. Non-invasive electrical and magnetic stimulation of the brain, spinal cord and roots: basic principles and procedures for routine clinical application. Report of an IFCN committee. Electroencephalogr Cinical Neurophysiol. 1994;91:79–92.

25. Dubois J, Kulikova S, Hertz-Pannier L, Mangin J-F, Dehaene-Lambertz G, Poupon C. Correction strategy for diffusion-weighted images corrupted with motion: application to the DTI evaluation of infants’ white matter. Magn Reson Imaging. 2014;32:981–992.

26. Rivière D, Geoffroy D, Denghien I, Souedet N, Cointepas Y. Anatomist: a python framework for interactive 3D visualization of neuroimaging data. Python Neurosci Work. 2011;:3–4.

27. Duclap D, Lebois LA, Schmitt WB, Riff O, Guevara P, Marrakchi-Kacem L, Schmitt WB. Connectomist-2.0: a novel diffusion analysis toolbox for BrainVISA. 2012.

28. Dobrynina L. Ischemic stroke in young adults: causes, clinical characteristics, diagnosis, prognosis for motor recovery. Dr. thesis. 2013. doi:10.1016/j.copsyc.2014.12.004.

29. Alt H, Godau M. Computing the Fréchet distance between two polygonal curves. Int J Comput Geom Appl. 1995;05:75–91.

30. Tunҫ B, Parker WA, Ingalhalikar M, Verma R. Automated tract extraction via atlas based adaptive clustering. Neuroimage. 2014;102:596–607.

31. Hayward KS, Neva JL, Mang CS, Peters S, Wadden KP, Ferris JK, Boyd LA. Interhemispheric Pathways Are Important for Motor Outcome in Individuals with Chronic and Severe Upper Limb Impairment Post Stroke. Neural Plast. 2017;2017:1–12.

32. Borich MR, Mang C, Boyd LA. Both projection and commissural pathways are disrupted in individuals with chronic stroke: investigating microstructural white matter correlates of motor recovery. BMC Neurosci. 2012;13:107.

33. Stewart JC, Dewanjee P, Tran G, Quinlan EB, Dodakian L, McKenzie A, See J, Cramer SC. Role of corpus callosum integrity in arm function differs based on motor severity after stroke. NeuroImage Clin. 2017;14:641–647.

34. Kemlin C, Moulton E, Lamy J-C, Houot M, Valabregue R, Leder S, Obadia MA, Meseguer E, Yger M, Brochard V, Corvol J-C, Samson Y, Rosso C. Elucidating the Structural and Functional Correlates of Upper-Limb Poststroke Motor Impairment. Stroke. 2019;50:3647–3649.

35. Witten IH, Frank E, Hall MA, Pal CJ. Data Mining, Fourth Edition: Practical Machine Learning Tools and Techniques, 4th ed. Morgan Kaufmann; 2016.

36. Perme MP, Manevski D. Confidence intervals for the Mann–Whitney test. Stat Methods Med Res. 2019;28:3755–3768.

37. Bonett DG, Wright TA. Sample size requirements for estimating pearson, kendall and spearman correlations. Psychometrika. 2000;65:23–28.

38. Lindenberg R, Zhu LL, Rüber T, Schlaug G. Predicting functional motor potential in chronic stroke patients using diffusion tensor imaging. Hum Brain Mapp. 2012;33:1040–1051.

39. Byblow WD, Stinear CM, Barber PA, Petoe MA, Ackerley SJ. Proportional recovery after stroke depends on corticomotor integrity. Ann Neurol. 2015;78:848–859.

40. Mang CS, Borich MR, Brodie SM, Brown KE, Snow NJ, Wadden KP, Boyd LA. Diffusion imaging and transcranial magnetic stimulation assessment of transcallosal pathways in chronic stroke. Clin Neurophysiol. 2015;126:1959–1971.

41. Domin M, Lotze M. Parcellation of motor cortex-associated regions in the human corpus callosum on the basis of Human Connectome Project data. Brain Struct Funct. 2019;224:1447–1455.

42. Senda J, Ito M, Watanabe H, Atsuta N, Kawai Y, Katsuno M, Tanaka F, Naganawa S, Fukatsu H, Sobue G. Correlation between pyramidal tract degeneration and widespread white matter involvement in amyotrophic lateral sclerosis: A study with tractography and diffusion-tensor imaging. Amyotroph Lateral Scler. 2009;10:288–294.

