## Supplementary materials for "Multimodal DTI-TMS assessment of the motor system in patients with chronic ischemic stroke"

#### Supplemental Methods

**** Spastic paresis scale of Institute of Neurology (1982) (Stoliarova, Kadykov, and Tkancheva 1982)**

- no movement problems – 0 points;
- light paresis (range of movements – 75-100% of norm; characteristics – slowness of movement) - 1 point;
- moderate paresis (range of movements – 50-75% of norm; characteristics – clumsy movements) – 2 points;
- apparent paresis (range of movements – 25-50% of norm; characteristics – global, synergistic) – 3 points;
- severe paresis (range of movements – 25% of norm; characteristics – very limited) – 4 points;
- plegia (lack of active movements) – 5 points.

#### Paired pulse TMS protocol

The paired pulse (ppTMS) was performed using the same stimulator and figure-of-eight monophasic coil in the contralesional hemisphere in all patients and in the ipsilesional hemisphere in all MEP+ patients. However, because RMT for monophasic TMS is higher than for biphasic TMS, in one out of 16 MEP+ patients, it was impossible to obtain MEPs using maximal intensity of the monophasic coil, and in two other patients, it was too painful. Thus, ppTMS of the ipsilesional side was performed only in 13 MEP+ patients. Two ppTMS paradigms, short-interval cortical inhibition (SICI) and intracortical facilitation (ICF), and single pulse paradigm were tested. For both ppTMS paradigms, intensities of 90% for the conditioning and 110% RMT for the test pulse were used [1]. The interstimulus intervals for the paired-pulse stimulation were 2 ms for SICI and 12 ms for ICF paradigms. SICI and ICF strength was calculated as a ratio: mean SICI amplitude/mean single pulse (SP) amplitude and mean ICF amplitude/mean SP amplitude.

### Supplementary Results

### Supplemental Tables

**Supplemental Table I.** Patients' demographic and infarction location characteristics for the three motor recovery outcome groups

| Motor recovery group | Gender | Age (y.o) | Brain infarct localization | Time after stroke (months) | DTI 1-yes, 0 -no | TMS 1-yes, 0 -no |
|---|---|---|---|---|---|---|
| I | male | 53 | subcortical | 52 | 1 | 1 |
| I | female | 55 | subcortical | 67 | 1 | 1 |
| I | female | 27 | subcortical | 14 | 1 | 1 |
| I | female | 33 | subcortical | 46 | 1 | 1 |
| I | male | 46 | subcortical | 12 | 1 | 1 |
| I | male | 54 | subcortical | 13 | 1 | 1 |
| I | male | 55 | cortico-subcortical | 20 | 1 | 1 |
| I | female | 57 | subcortical | 37 | 0 | 1 |
| I | female | 39 | cortico-subcortical | 46 | 0 | 1 |
| I | female | 37 | cortico-subcortical | 25 | 1 | 0 |
| | | | | | 8 | 9 |
| II | female | 46 | cortico-subcortical | 30 | 1 | 1 |
| II | male | 48 | subcortical | 21 | 1 | 1 |
| II | male | 52 | subcortical | 37 | 1 | 1 |
| II | male | 48 | cortico-subcortical | 25 | 1 | 1 |
| II | male | 52 | subcortical | 99 | 1 | 1 |
| II | male | 58 | cortical | 26 | 1 | 1 |
| II | male | 65 | cortico-subcortical | 15 | 0 | 1 |
| | | | | | 6 | 7 |
| III | male | 57 | cortico-subcortical | 12 | 1 | 1 |
| III | female | 36 | cortico-subcortical | 22 | 1 | 1 |
| III | male | 49 | subcortical | 67 | 1 | 0 |
| III | female | 40 | subcortical | 14 | 1 | 1 |
| III | female | 40 | cortico-subcortical | 16 | 1 | 1 |
| III | male | 50 | cortico-subcortical | 38 | 1 | 1 |
| III | male | 58 | subcortical | 6 | 1 | 1 |
| III | female | 49 | subcortical | 29 | 1 | 1 |
| III | male | 58 | cortico-subcortical | 67 | 1 | 1 |
| III | male | 44 | cortico-subcortical | 14 | 1 | 1 |
| III | female | 41 | cortico-subcortical | 58 | 1 | 1 |
| III | female | 26 | subcortical | 7 | 1 | 1 |
| III | female | 41 | subcortical | 10 | 1 | 1 |
| III | male | 37 | cortico-subcortical | 7 | 1 | 1 |
| III | male | 66 | subcortical | 6 | 1 | 1 |
| III | male | 35 | cortico-subcortical | 26 | 1 | 1 |
| III | male | 43 | cortico-subcortical | 14 | 1 | 0 |
| III | female | 36 | cortico-subcortical | 21 | 1 | 0 |
| | | | | | 18 | 15 |

**Supplemental Figures**

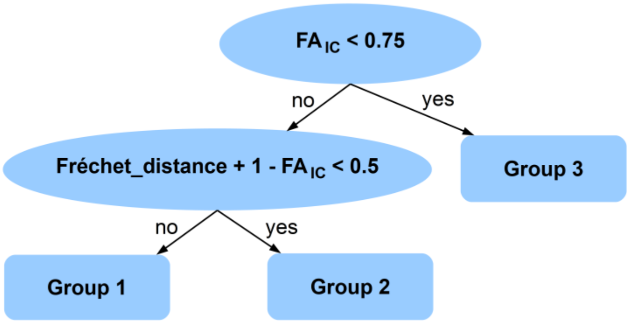


**Supplemental Figure I:** Decision tree for 3 class patients' classification using FA measures in the CST. While FA asymmetry in IC perfectly separates Groups 1 and 2 from Group 3 ($FA_{IC}$ < 0.75), a linear combination of Fréchet distance between FA profiles, from contra- and ipsilesional hemispheres, and FA in IC (Fréchet distance +(1-$FA_{IC}$) < 0.5), makes it possible to separate all three motor recovery groups from each other with 91% accuracy.

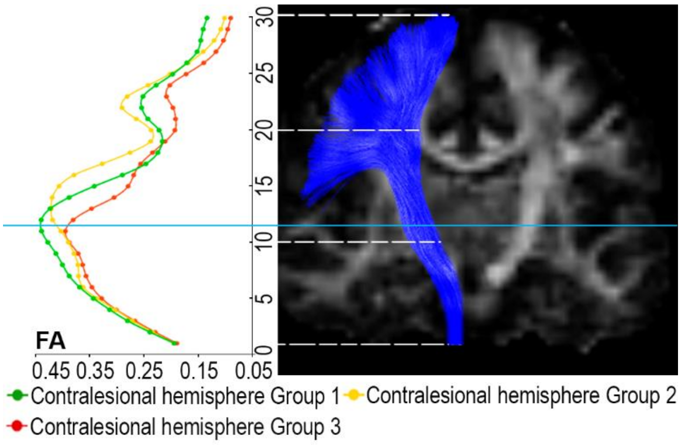


**Supplemental Figure II**: FA CST profiles for the contralesional hemisphere for all three groups. Light blue line is reflecting the level of IC.

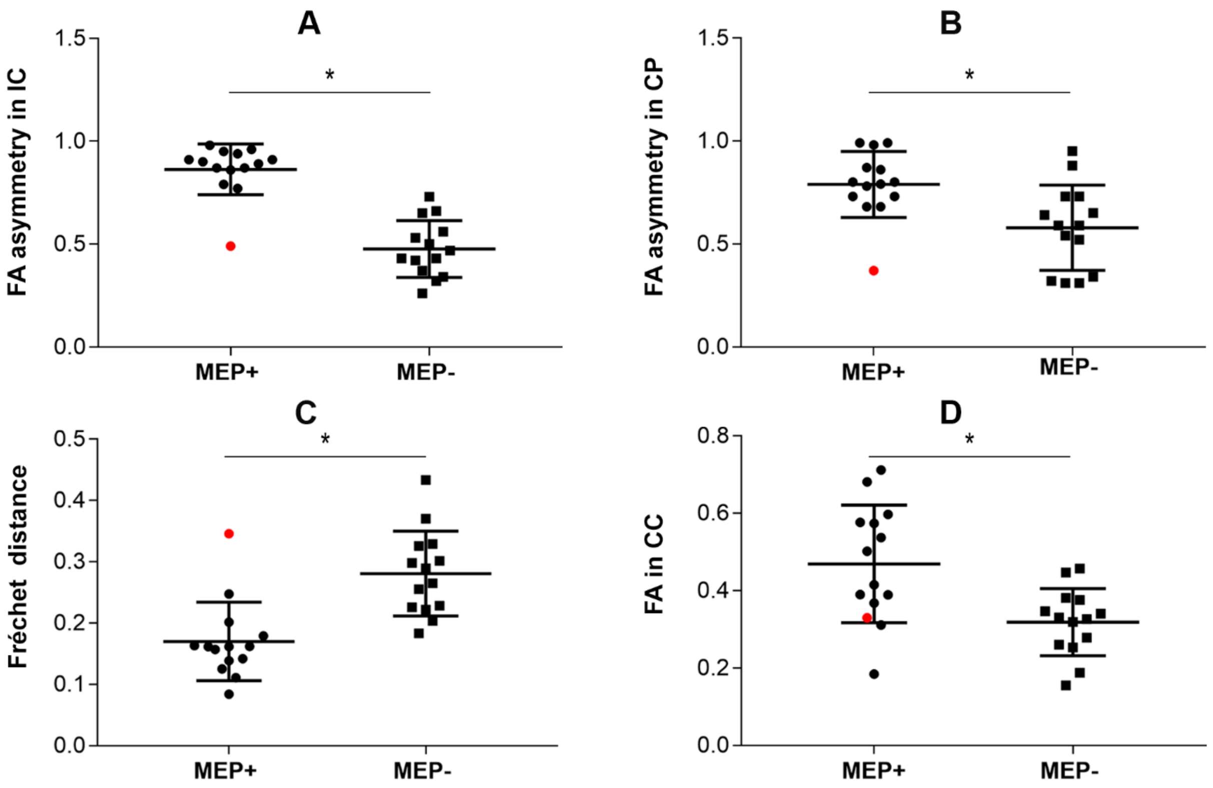


**Supplemental Figure III:** Difference of CST and corpus callosum (CC) structural integrity between MEP+ and MEP- patients. (A) Difference of fractional anisotropy (FA) ratio in the internal capsule (IC). One can see that MEP+ and MEP- groups do not overlap except for the only outlier (this outlier patient is highlighted with red in all the graphs) (Mann-Whitney U-test: *θ = 0.97 [95% CI, 0.78 to 1.00], *p< 0.0001); (B) Difference of FA ratio in the cerebral peduncle (CP) (Mann-Whitney U-test: *θ = 0.82 [95% CI, 0.58 to 0.93], *p =0.0046) (C) Difference of Fréchet distance between ipsi- and contralesional CST FA profiles (Mann-Whitney U-test: *θ = 0.91 [95% CI, 0.67 to 1.00], *p = 0.0003) (D) Difference of FA in the CC (Mann-Whitney U-test: *θ = 0.80 [95% CI, 0.57 to 0.92], *p = 0.0072).

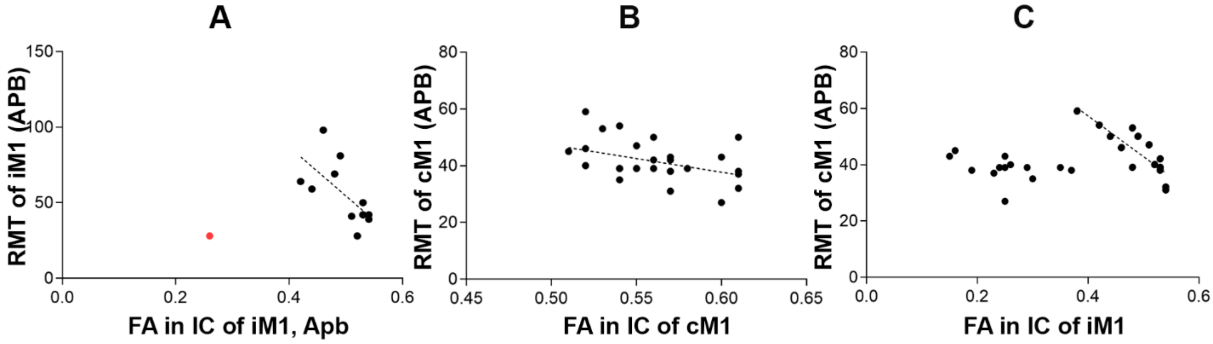


**Supplemental Figure IV:** Spearman correlation between RMTs and IC FA (A) ipsilesional IC FA and RMT correlated highly if we do not consider the outlier from the bad recovery group with very low ipsilesional RMTs (Spearman correlation coefficient = -0.66, p=0.0233, [95% CI, -0.88 to -0.19]); (B) contralesional IC FA and RMT showed smaller but more significant linear correlation (Spearman correlation coefficient= -0.46, p=0.0143, [95% CI, -0.71 to -0.10]); (C) while correlating contralesional RMT with ipsilesional IC FA values, patients could be divided into two clusters: in patients with higher IC FA (FA>0.4) it correlated strongly and negatively with RMT (Spearman correlation coefficient= -0.86, p=0.0002, [95% CI, -0.95 to -0.63]), while in patients with lower IC (FA<0.4), no correlation was observed.

### Supplementary discussion

#### Excitation/inhibition balance of the contralesional motor cortex probed by ppTMS

Our main question, concerning the CC integrity, was whether it is associated with the excitation/inhibition balance of the contralesional hemisphere. Despite the fact, that brain stimulation approaches, dedicated to change the excitation/inhibition balance between hemispheres, have been used for more than 10 years [2], the actual state of the contralesional hemisphere and its dependence on the interhemispheric connections, is still not fully understood. Only a few previous studies have examined the relationship between contralesional motor cortex excitability and ppTMS phenomena, along with the interhemispheric inhibition, to address the question whether abnormally increased contralesional primary motor cortex excitability results in an excessive inhibition of the contralesional motor cortex [3,4]. We have not found studies where ppTMS phenomena in the contralesional hemisphere were interrelated with the CC integrity. In our sample we did not see any correlation between the CC integrity and the contralesional motor cortex functional state estimated with TMS. At the same time, we observed no disinhibition or excitation of the contralesional motor cortex in the bad recovery outcome group compared to other groups. Conversely, lack of SICI phenomena in the contralesional motor cortex occurred in some moderately and well recovered patients. Thus, our results are in agreement with a bimodal-balance recovery model, proposing to inhibit excessively hyperexcited contralesional hemisphere only in less severely affected patients [5].

### Supplemental references


1. Lioumis P, Mustanoja S, Bikmullina R, Vitikainen A-M, Kičić D, Salonen O, Tatlisumak T, Kaste M, Forss N, Mäkelä JP. Probing modifications of cortical excitability during stroke recovery with navigated transcranial magnetic stimulation. *Top Stroke Rehabil*. 2012;19:182–192.

2. Ovadia-Caro S, Khalil AA, Sehm B, Villringer A, Nikulin V V., Nazarova M. Predicting the response to non-invasive brain stimulation in stroke. *Front Neurol*. 2019;in press:302.

3. Bütefisch CM, Weβling M, Netz J, Seitz RJ, Hömberg V. Relationship Between Interhemispheric Inhibition and Motor Cortex Excitability in Subacute Stroke Patients. *Neurorehabil Neural Repair*. 2008;22:4–21.

4. Guggisberg AG, Koch PJ, Hummel FC, Buetefisch CM. Brain networks and their relevance for stroke rehabilitation. *Clin Neurophysiol*. 2019;130:1098–1124.

5. Pino G Di, Pellegrino G, Assenza G, Capone F, Ferreri F, Formica D, Ranieri F, Tombini M, Ziemann U, Rothwell JC. Modulation of brain plasticity in stroke : a novel model for neurorehabilitation. *Nat Publ Gr*. 2014;10:597–608.
